# Functional characterization of the cnidarian antiviral immune response reveals ancestral complexity

**DOI:** 10.1101/2020.11.12.379735

**Authors:** Magda Lewandowska, Ton Sharoni, Yael Admoni, Reuven Aharoni, Yehu Moran

## Abstract

Animals developed a broad repertoire of innate immune sensors and downstream effector cascades for defense against RNA viruses. Yet, this system highly varies between different bilaterian animals, masking its ancestral state. In this study we aimed to characterize the antiviral immune response of the cnidarian *Nematostella vectensis* and decipher the function of the retinoic acid-inducible gene I-like receptors (RLRs) known to detect viral double-stranded RNA (dsRNA) in bilaterians, but activate different antiviral pathways in vertebrates and nematodes. We show that a mimic of long viral dsRNA triggers a complex antiviral immune response bearing features distinctive for both vertebrate and invertebrate systems. Furthermore, the results of affinity assays and knockdown experiments provide functional evidence for the conserved role of RLRs in initiating immune response to dsRNA that originated before the cnidarian-bilaterian split and lay a strong foundation for future research on the evolution of the immune responses to RNA viruses.

## Introduction

The immune system has long been known for its remarkable patterns of rapid evolution owing to strong selective drivers such as fast-evolving pathogens^1,2^. Thus, revealing conservation among phylogenetically distant lineages can provide unprecedented insights into the evolution of these defense mechanisms. For instance, it has been recently reported that both eukaryotic antiviral DNA-sensing mechanism driven by cGAS-STING axis and the downstream inhibitors of virus replication called viperins have originated in procaryotes as anti-bacteriophage mechanisms^3–5^. Viruses are very often sensed by their nucleic acids which bear features not shared by their host cells^6,7^. Specifically, eukaryotes had to adapt to emerging RNA viruses by developing strategies to recognize such non-self genetic material. The best characterized foreign features are i) double-stranded RNA (dsRNA) structures and ii) triphosphate on 5’ ends, both of which are mostly absent during host cell homeostasis but are accumulated in viral infection, either directly derived from the viral genomes or formed as the replication or transcription intermediates^8–10^. In plants, nematodes and arthropods, the presence of the cytoplasmatic dsRNA triggers RNA interference (RNAi) which involves slicing dsRNA into short interfering RNAs (siRNA) by the ribonuclease III Dicer, often followed by signal amplification by RNA-dependent RNA polymerases (RdRPs) and final silencing of viral RNA by Argonaute proteins^11–14^. In vertebrates, dsRNA is detected by several families of pattern-recognition receptors (PRRs) which trigger downstream expression of type I interferons (IFNs) and other proinflammatory cytokines^7^. Retinoic acid-inducible gene I (RIG-I) -like receptors (RLRs) is a family of metazoan-specific ATP-dependent DExD/H box RNA helicases that function as the major cytoplasmic PRRs binding dsRNA^15–17^ (**Fig. 1a**). In vertebrates, ligands of RIG-I and its paralog melanoma differentiation-associated protein 5 (MDA5) include short, blunt-end dsRNA with 5’ di- and triphosphate^9,18–22^ and long irregular dsRNA^23–26^, respectively. Caspase activation and recruitment domains (CARDs) of RIG-I and MDA5 (**Fig. 1b**) are necessary for regulation, oligomerization and subsequent interaction with adaptor molecules to trigger downstream effector cascades^27^. Absence of the CARD domain in the third vertebrate RLR – Laboratory of Genetics and Physiology 2 (LGP2) – prevents signal transduction and is correlated with its dual regulatory functions^28^.

**Figure 1.**
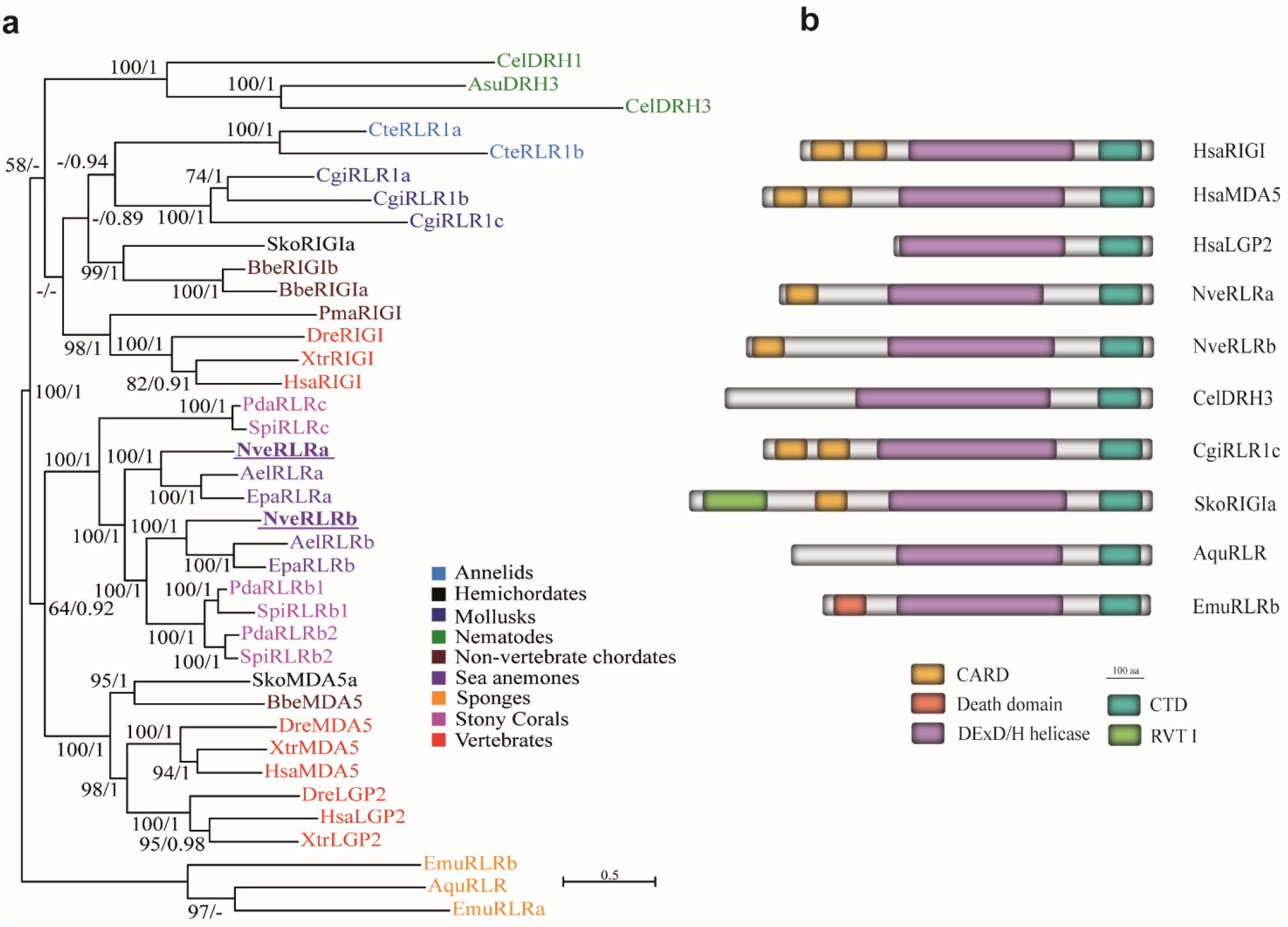
Phylogenetic relationship of metazoan RLRs. **(a)** Maximum likelihood and Bayesian inference consensus phylogenetic tree of representative RLR sequences, bootstrap values above 50% are presented for each node. Posterior probability values of a Bayesian tree of the same topology between 0.85-1.0 are indicated for each node. Ael, *Anthopleura elegantissima*, Aqu, *Amphimedon queenslandica*, Asu, *Ascaris suum*, Bbe, *Branchiostoma belcheri*, Cte, *Capitella teleta*, Cgi, *Crassostrea gigas*, Cel, *Caenorhabditis elegans*, Dre, *Danio rerio*, Emu, *Ephydatia muelleri*, Epa, *Exaiptasia pallida*, Has, *Homo sapiens*, Nve, *Nematostella vectensis*, Pda, *Pocillopora damicronis*, Pma, *Petromyzon marinus*, Sti, *Stylophora pistillata*, Xtr, *Xenopus tropicalis*. **(b)** Schematic representation of selected RLR representatives of major phylogenetic groups. CARD - caspase recruitment domain; CTD - C-terminal domain; RVT I – reverse transcriptase.

Although RLRs have been found in many animal phyla (**Fig. 1a**) and display structural conservation **(Fig. 1b)**, their function in invertebrate immune response remains understudied in animals other than vertebrates and nematodes, leaving a major gap in the understanding of RLRs evolution. In this study we aimed to characterize the immune response to viral dsRNA mimics in *Nematostella vectensis*, a model organism of phylum Cnidaria (sea anemones, corals, jellyfish and hydroids) separated from its sister group Bilateria (the majority of extant animals, including vertebrates and nematodes) by > 600 million years of evolution^29,30^. We observe in this cnidarian a strong immune response triggered by long, but not short 5’ triphosphate-bearing dsRNA which supports our phylogenetic analyses of RLRs. We show that both *N. vectensis* RLRs (NveRLRs) are likely to take part in the antiviral immune response and that one of them is showing affinity to long dsRNA. Finally, knockdown of this RLR results in impaired downstream effector processes suggesting its key role in initiating immune response to dsRNA.

## Results

### Ancestral RLRs duplication likely predates the Bilateria-Cnidaria split

In order to gain a better understanding of the evolutionary fate of RLRs and the position of *N. vectensis* homologs within the family of these viral nucleic acid sensors, we reconstructed previous phylogenetic trees with an addition of numerous recently available sequences. Instead of including other distantly related DExD/H helicases, such as RNA-specific endoribonuclease Dicer or elongation initiation factor 4A (eIF4A), we performed phylogenetic analysis exclusively of RLRs with the sequences of sponges, one of the first two metazoan phyla to diverge^31,32^, set as an outgroup (**Fig. 1a**). Similar to previous studies^33,34^, we have not identified any RLRs homologs in Placozoa and Ctenophora. Within Cnidaria, we identified RLRs sequences in Hexacorallia (sea anemones and stony corals) while they are absent in the Meduzosoa clade, clearly indicating a loss. Interestingly, unlike in previous studies^33,34^, we observed a well-supported clustering of all hexacorallian RLRs within bilaterian MDA5/LGP2 clade, which forms a sister group to bilaterian RIG-I sequences. This unexpected finding suggests that, in contrast to the previous hypothesis^35^, RLRs paralogs duplicated before the split of Bilateria and Cnidaria and all cnidarian RLRs paralogs originated from a MDA5/LGP2 ancestral protein. Furthermore, both *Nematostella* CARD-containing protein sequences – NveRLRa and NveRLRb – are positioned in separated clades with orthologs from other sea anemones suggesting their ancient duplication predating sea anemone divergence and most likely the functional non-redundancy. Clustering of RLRb sequences of sea anemones within one of the clades of stony corals which have split 320 million years ago^36^ further supports the hypothesis of an ancient sub- or neofunctionalization of the sea anemone RLRs.

### Lack of response in *Nematostella* to RIG-I specific ligand

To functionally support our observation that *Nematostella* RLRs are closer related to Bilateria MDA5 receptor, we decided to first employ known ligand affinity and test *Nematostella* response to MDA5 and RIG-I-specific ligands. To this end, we microinjected *N. vectensis* embryos with polyinosinic:polycytidylic acid (poly(I:C)), a mimic of long viral dsRNA and a potent agonist of MDA5^25,26^, and a short dsRNA 19-mer with 5’ triphosphate group (5’ppp-dsRNA) which is known to be detected by RIG-I^9,19^. Analysis of differentially expressed genes (DEG) upon the treatments with viral mimics revealed a strong response to poly(I:C) (**Fig. 2a, Supplementary File S1, Supplementary Fig. S1d,e**) with a peak of the differential expression at 24 hours post-injection (hpi) accounting for 67.26% of variance revealed by Principal Component 1 (**Fig. 3c)**. Among 3 different time points, we have observed an almost complete lack of transcriptomic response in 6 hpi (*n* of DEG = 14) which agrees with a low transcript abundance at the onset of zygotic transcription in *Nematostella*^37^. Both at 24 and 48 hpi (*n* of DEG = 1475 and 524, respectively) the majority of DEG were upregulated (**Fig. 3a,b**) which is a common pattern of the innate immune response to viral ligands^38^. In contrast, the transcriptomic response to vertebrate RIG-I specific dsRNA ligand revealed a striking lack of signature of the antiviral immune processes (**Fig. 2b, Supplementary File 1, Supplementary Fig. S1a,b,c**) despite being applied at 90 – 180-fold higher concentration compared to concentrations used for vertebrates, suggesting that unlike in vertebrates^39–41^, a triphosphate group on 5’ blunt-end of short dsRNA is not triggering an immune reaction in *N. vectensis*.

**Figure 2.**
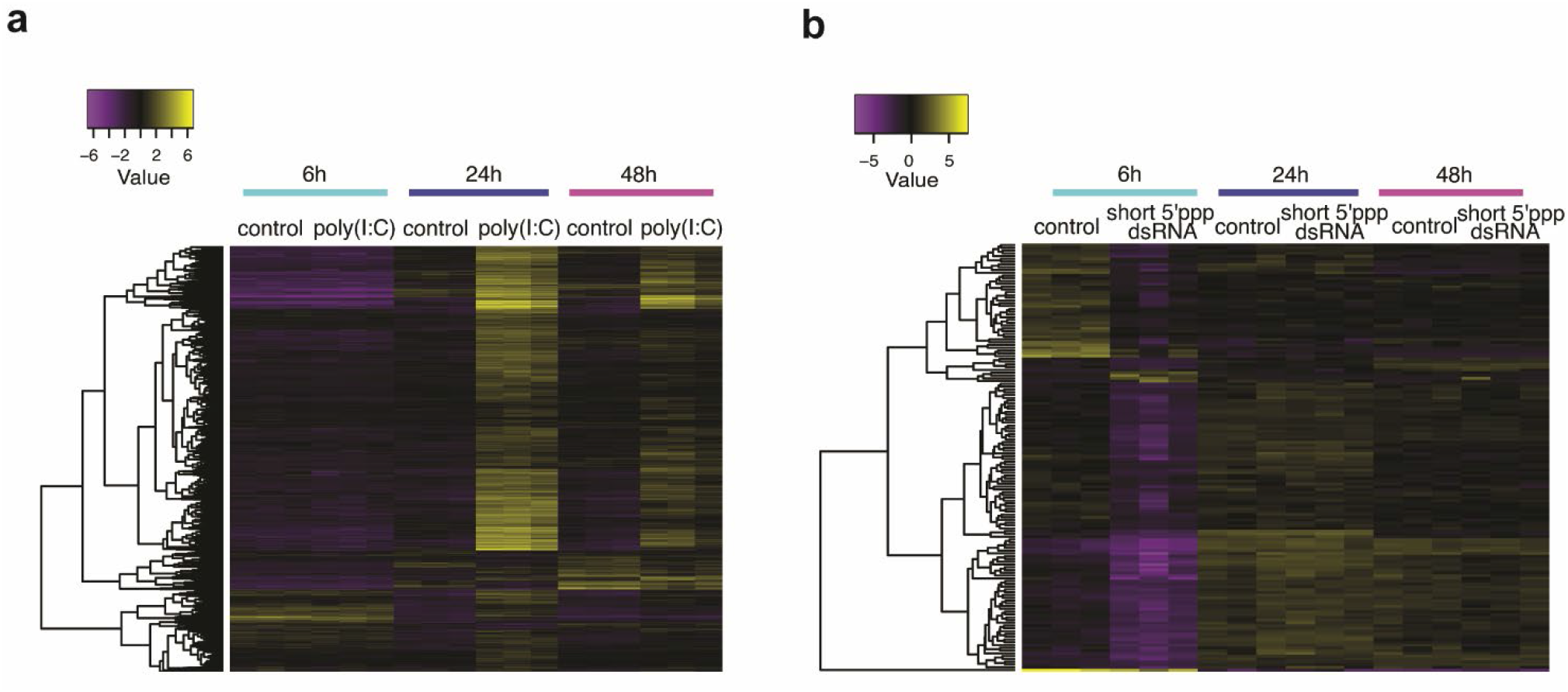
Differential gene expression after microinjections of viral mimics. Heatmap of differentially expressed genes upon administration of **(a)** poly(I:C) vs 0.9% NaCl serving as a control, and **(b)** 19-mer dsRNA with 5’ triphosphate and 19-mer dsRNA with 5’ hydroxyl group serving as a control.

**Figure 3.**
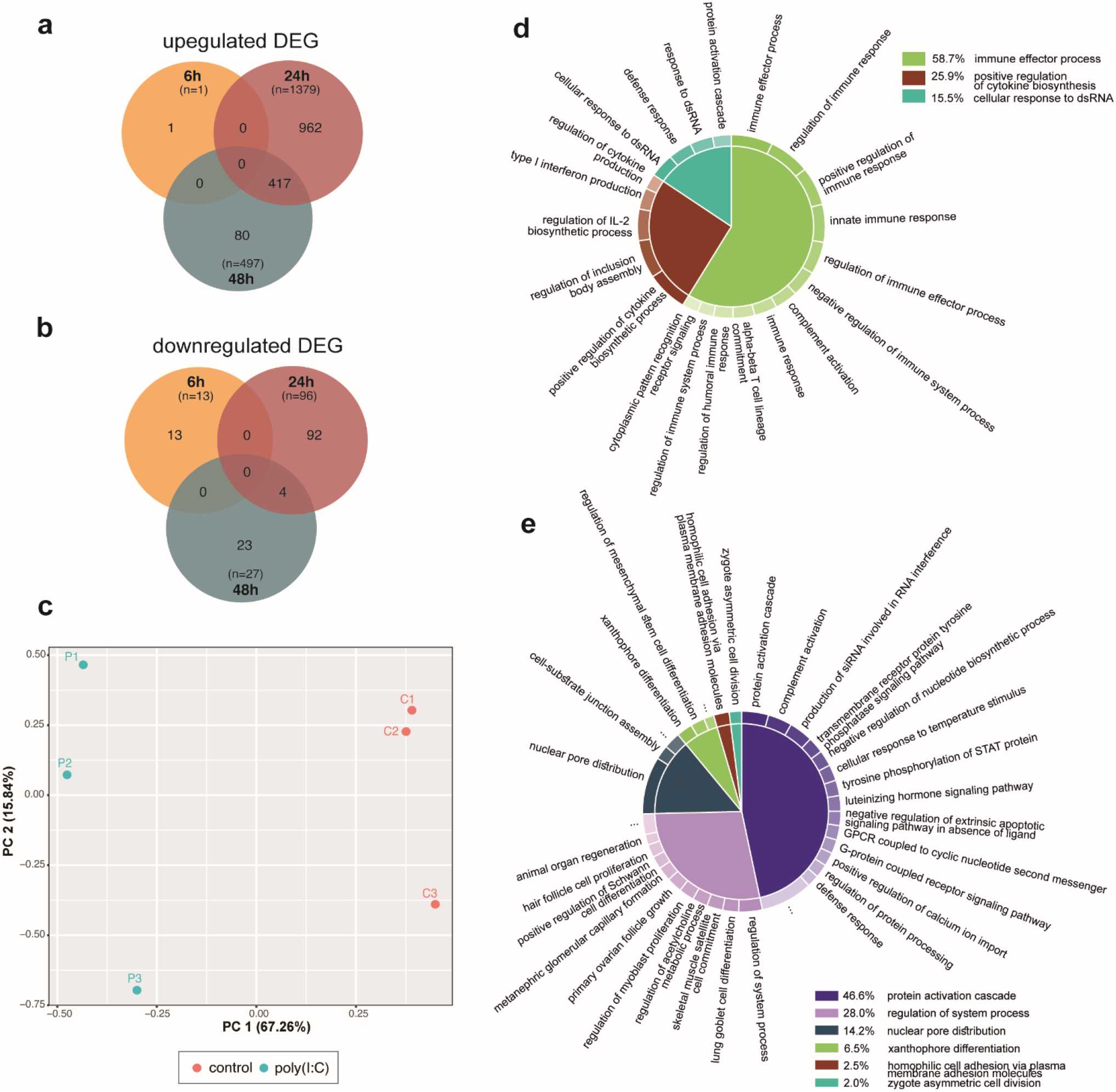
Signature of the innate immune response to poly(I:C). Venn diagram of differentially expressed genes which were **(a)** upregulated and **(b)** downregulated after poly(I:C) administration. **(c)** PCA plot representing whole transcriptome of poly(I:C)-injected animals at 24 hpi. **(d)** GO terms enrichment results of DEG upregulated at 24 hpi and **(e)** 48 hpi.

Results of the gene-set enrichment analysis (GSEA) revealed the abundance of gene ontology (GO) terms related to the innate immunity and strengthened our inference on strong antiviral response triggered by poly(I:C) at 24 hpi (**Fig. 3d, Supplementary File S2**) and to a lesser extent at 48 hpi (**Fig. 3e, Supplementary File S2**). Importantly, the vast majority of responding genes at the later stage overlaps with the upregulated genes of the former one (**Fig. 3a**), suggesting a continuous attenuating immune response. In all tested groups enriched GO terms contained many vertebrate-specific terms, therefore we had to treat it as an approximation to a true gene function. Although the GSEA for the short 5’ppp-dsRNA had not revealed enriched GO terms which would pass the statistical threshold likely due to the low DEG abundance (**Supplementary File S2**), we decided to examine the only DEG group responding to this treatment i.e. genes downregulated at 6 hpi (**Fig. 2b**). Identified GO terms groups were predominantly related to the early-stage development (**Supplementary Fig. S1f**) which led us to the hypothesis that the presence of very high molarity of charged compounds might either directly or indirectly interfere with the onset of zygotic transcription, possibly by altering the cellular pH or disrupting physiological processes through the divalent cations chelating activity^42^.

### Response to poly(I:C) reveals patterns of both invertebrate and vertebrate antiviral innate immunity

Among poly(I:C)-upregulated genes at 24 hpi we identified both of *Nematostella* RLRs, with more significant increase for *NveRLRb* (edgeR-based log_2_ Fold Change (FC) = 3.275, False Discovery Rate (FDR) = 1.54e-20) than for *NveRLRa* (score below the fold change threshold, i.e. edgeR-based log_2_FC = 1.864, FDR = 6.55e-09). This increase suggests a possibility of a feed-forward loop similar to that observed in vertebrate antiviral immune response^43^. Moreover, many genes linked to RNAi (e.g. *NveDicer1*, *NveAGO2*, *NveRdRP1-3*) and numerous homologs of genes involved in antiviral innate immune response in both vertebrates and invertebrates animals^7,44^ (e.g. Interferon regulatory factors (IRFs), RNAse L, guanylate-binding proteins (GBPs), 2’-5’-oligoadenylate synthetase 1 (OAS1), nuclear factor kappa-light-chain-enhancer of activated B cells (NF-κB), radical SAM domain-containing 2 (Viperin), to mention few, **Supplementary File S1**) were also detected. Interestingly, we observed a significant upregulation from a previously undescribed factor (gene symbol: NVE23912) which is a cysteine-rich sequence (11 cysteine residues) with a predicted signal peptide and no significant homology to any known genes. Our search for homologs in Transcriptome Shotgun Assembly (TSA) and NCBI nr databases revealed that it is likely a secreted hexacorallian-specific protein (**Supplementary Fig. S2**) which resembles pattern of proteins under strong selective pressure displayed by the high conservation limited to the cysteine positions. Altogether, the described features make it a good candidate for further functional studies which could validate whether this novel factor is playing an important role in the innate immunity of *N. vectensis* and possibly other members of Hexacorallia.

To get wider view of the nature of poly(I:C)-induced DEG we examined promoter sequences of the induced genes by two different approaches. First, we screened the coding strand for the presence of the TATA-box in both the close proximity to the transcription starting site (TSS) (38 bp) and in a more permissive screening window (100 bp upstream and 100 bp downstream of TSS). It has been previously suggested that mammalian immune-related genes which are rapidly diverging and exhibit greater levels of expression variability across individual cells, such as cytokines and chemokines, share a common promoter architecture enriched in TATA-boxes^45^. Interestingly, *N. vectensis* displays a significant increase in abundance of TATA-box elements in poly(I:C)-upregulated genes when searching both window sizes which seems to correlate with the level of genes inducibility (**Supplementary Fig. S3a,b, Supplementary File S3**). Within protein sequences of TATA-box containing genes, we predicted a similar enrichment of signal peptides suggesting that many of these proteins might be involved in secretory pathways (**Supplementary Fig. S3c,d, Supplementary file S3**). Furthermore, the search of known transcription factor binding sites (TFBS) revealed numerous motifs known to be involved in regulating transcription of antiviral immune-related genes in vertebrates such as those recognized by STATs, IRFs, NF-κB or members of ETS family^46–49^ (**Supplementary File S3**). In order to circumvent the limitation of using the vertebrate motif matrix, we scanned the *N. vectensis* genome for the presence of the homologs of vertebrate immune-related transcription factors. Importantly, we have identified numerous candidate homologs of these factors in *N. vectensis* genome among which a large group showed upregulation in response to poly(I:C) treatment supporting the notion that they might play role in orchestrating the observed immune response (**Supplementary File S3**).

### Role of NveRLRs in detecting long dsRNA

To confirm the results of our RNA-seq DEG analysis we assayed gene expression in independent biological replicates. RT-qPCR analysis at 24 hpi validated the upregulation of *NveRLR*s (relative expression_*NveRLRa*_ = 1.98, 95% CI, 1.042 – 3.494, p-value = 0.0425, relative expression_*NveRLRb*_ = 5.795, 95% CI, 3.992 – 8.411, p-value = 0.000643) (**Fig. 4a**), as well as several other putative immune-related genes (**Supplementary Fig. S4d, Supplementary File S4**) in response to poly(I:C) treatment, and an unaffected expression level of *NveRLR*s transcripts when treated with short 5’ppp-dsRNA (relative expression_*NveRLRa*_ = 0.815, 95% CI, 0.468 –1.420, p-value = 0.325, relative expression_*NveRLRb*_ = 1.071, 95% CI, 0.528 – 2.173, p-value = 0.778) (**Fig. 4b**). Importantly, the examination of the *NveRLR*s mRNA levels in response to the control treatments did not reveal a significant background upregulation which could distort the results of ligand specificity (**Supplementary Fig. 5a,b**). To confirm these results at the protein level, we generated custom polyclonal antibodies against *N. vectensis* RLRs which specificity has been tested beforehand. NveRLRs levels were tested at 48 hpi in order to diminish the effect of maternally deposited proteins. The result of Western blot confirmed strong upregulation of both NveRLRs after poly(I:C) stimulation (**Fig. 4c,d**) which correlates with the increased transcript abundance.

**Figure 4.**
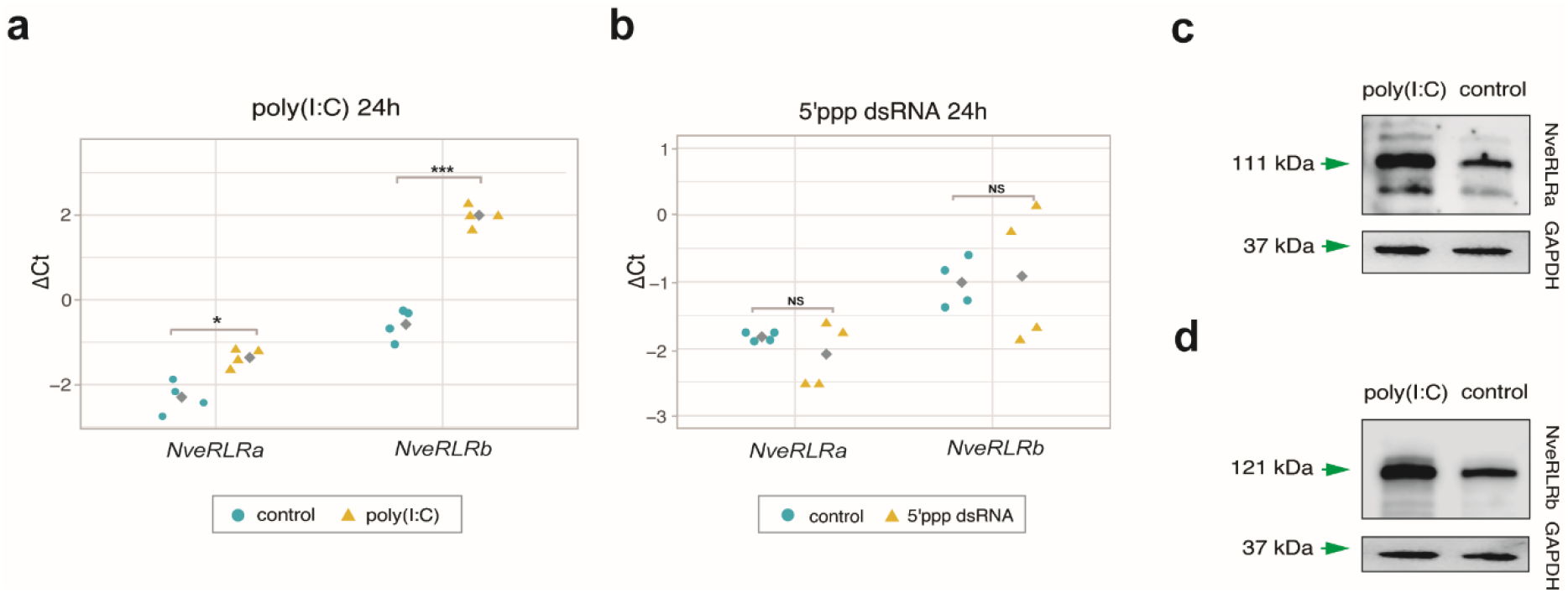
Response of *Nematostella* putative dsRNA helicases to viral mimics. *NveRLR*s mRNA expression level measured by RT-qPCR in response to **(a)** poly(I:C) and **(b)** short 5’ ppp dsRNA. Grey squares represent mean values. Western blot validation of **(c)** NveRLRa and **(d)** NveRLRb protein level in response to poly(I:C) at 48 hpi. Significance level was assessed by paired two-tailed Student’s t-test; * p value < 0.05, ** p value <0.01, *** p value <0.001, NS – not significant.

Next, we aimed to examine the ability of NveRLRs to bind poly(I:C). To this end, we generated two *N. vectensis* transgenic lines, each expressing FLAG-tagged *NveRLR* and a fluorescent mCherry gene under a ubiquitous promoter of the TATA-Box Binding Protein (TBP) gene (**Fig. 5a**). Progeny of F_1_ female heterozygotes and wild-type animals was collected directly after fertilization (0 h) and the presence of maternally deposited FLAG-tagged RLRs was confirmed (**Fig. 5b**). In vitro binding assays of poly(I:C) covalently linked to biotin on wild-type protein extracts confirmed specificity of mouse FLAG antibody (**Fig. 5c**). The results of the in vitro poly(I:C) binding on the transgenic lines revealed a significant enrichment of NveRLRb in poly(I:C)-biotin pulldown samples indicating specific binding of long dsRNA by NveRLRb (**Fig. 5c**). Unexpectedly, no poly(I:C) affinity was detected when assaying NveRLRa (**Fig. 5d**). In order to monitor how accurately the conditions of transgenic expression mimic the native proteome composition, we examined levels of NveRLRs levels in recently published mass spectrometry data spanning different developmental stages of *N. vectensis*^50^. Interestingly, we noticed that while NveRLRb displays relatively stable expression throughout the lifecycle, levels of NveRLRa in the unfertilized egg are below the detection threshold (in agreement with previous proteomic studies of *Nematostella* eggs^51,52^) and show significantly lower expression than NveRLRb across all developmental stages (**Supplementary Fig. S6**). It is therefore plausible that NveRLRa carries regulatory function or binds yet uncharacterized ligands or, alternatively, that its ligand specificity matures along with the development due to co-expression of other crucial factors.

**Figure 5.**
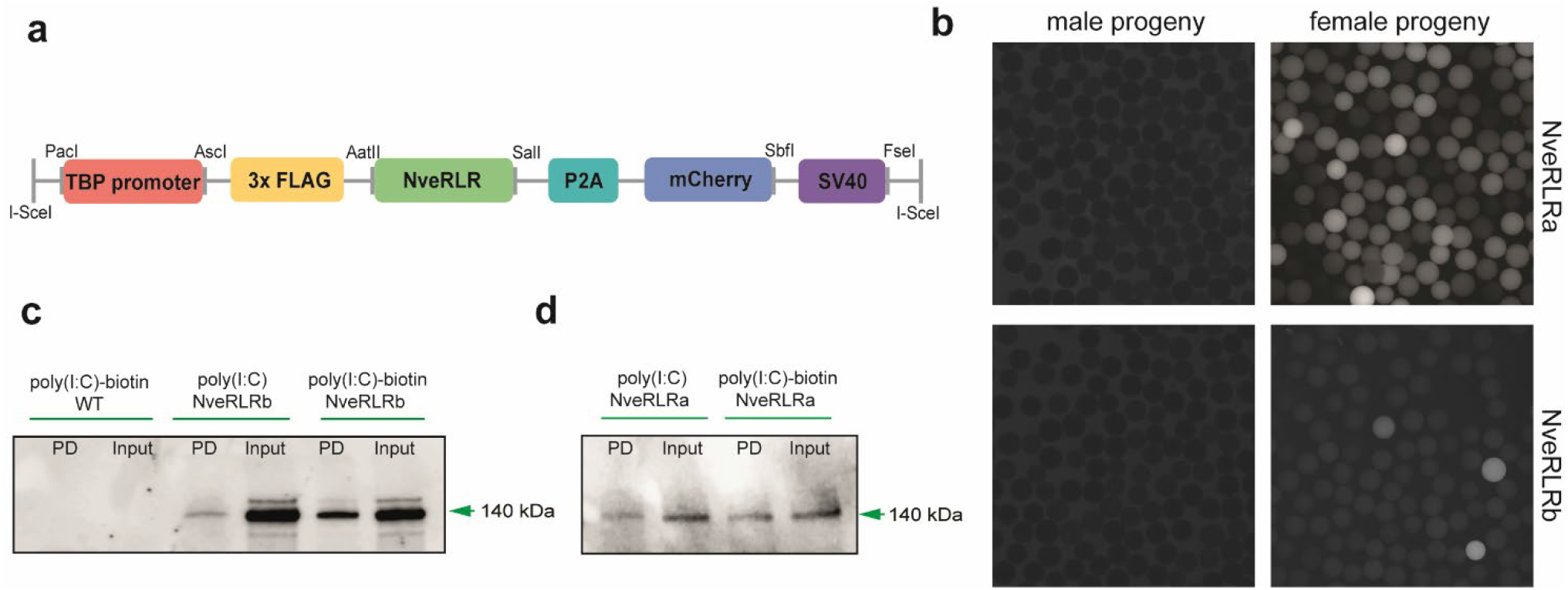
NveRLRs affinity to poly(I:C). **(a)** Schematic representation of the FLAG-*NveRLR* construct (7,071 bp and 7,643 bp for *NveRLRa* and *NveRLRb*, respectively) used for transgenesis. TBP promoter, self-cleaving P2A sequence, mCherry gene and polyadenylation signal SV40 are also shown. **(b)** Maternal deposition of the FLAG-NveRLR observed after crossing transgenic females (right panels) with WT males; fluorescent protein is missing in transgenic male progeny (left panels). **(c)** Results of poly(I:C)-biotin *in vitro* binding assay showing affinity of FLAG-NveRLRb but not **(d)** FLAG-NveRLRa to poly(I:C); PD – pulldown.

### Knockdown of *NveRLRb* interferes with the *in vivo* response to long dsRNA

Poly(I:C)-induced upregulation of *NveRLR*s both at the gene and the protein levels and FLAG:NveRLRb affinity to poly(I:C)-biotin led us to the assumption that both proteins might carry an important function in detecting viral dsRNA and hence, orchestrating downstream antiviral immune processes in *Nematostella.* To further corroborate this theory, we generated knockdown (KD) animals by microinjection of short hairpin RNA (shRNA) targeting 3 different regions of each of *NveRLR*s. The initial validation assays of KD efficiency and shRNAs immunogenicity revealed a strong (~85-90%) and moderate (~60%) effect of all *NveRLRb* and *NveRLRa* shRNAs, respectively (**Supplementary Fig. S5c,d**), and very low impact on the expression levels of putative immune-related genes of all shRNAs (**Supplementary Fig. S5e-j**). Due to the lack of strong knockdown effect by all candidate *NveRLRa* shRNAs, we decided to include all 3 tested variants for this gene and 2 shRNAs for *NveRLRb*. Following the assumption that NveRLRs might act as sensors in antiviral immune response, we co-injected each shRNAs with poly(I:C) and tested at 48 hpi the mRNA levels of candidate genes previously proved to respond to the poly(I:C) treatment. We classified genes as those affected by *NveRLR*s KD when their expression was significantly different when compared to control shRNA and obtained by at least two different shRNAs. The first unexpected observation was that while *NveRLRb* KD efficiency remained comparable to the initial screening assays (~90%), *NveRLRa* KD level decreased to approximately 45% (**Fig. 6a,b, Supplementary File S4**). Of note, none of the *NveRLR*s KD experiments exerted a strong and ubiquitous reciprocal effect on the other sensor. Importantly, knockdown of *NveRLRb* resulted in noticeable downregulation of both tested components of RNAi i.e. *NveDicer1* (relative expression_shRNA1_ = 0.370, 95% CI, 0.242 – 0.565, p-value = 0.005; relative expression_shRNA2_ = 0.165, 95% CI, 0.106 – 0.257, p-value = 0.001) and *NveAGO2* (relative expression_shRNA1_ = 0.663, 95% CI, 0.53 – 0.83, p-value = 0.01; relative expression_shRNA2_ = 0.451, 95% CI, 0.272 – 0.547, p-value = 0.00095), as well as *NveIRF1* (relative expression_shRNA1_ = 0.401, 95% CI, 0.236 – 0.683, p-value = 0.012; relative expression_shRNA2_ = 0.262, 95% CI, 0.171 – 0.402, p-value = 0.00215) (**Fig. 6c,d,e, Supplementary File S4**) and an apparent but not significant decrease in expression of hexacorallian-specific factor NVE23912 (**Supplementary Fig. S4c, Supplementary File S4**). Interestingly, neither *NveOAS1* nor *NveGBP1* mRNA levels were significantly affected by the *NveRLRb* shRNA-poly(I:C) co-injection (**Supplementary Fig. S4a,b, Supplementary File S4**). In contrast to *NveRLRb* KD, response to *NveRLRa* shRNAs did not reveal any clear signature of the impaired downstream process in 5 out of 6 tested genes and displayed a general pattern of high expression variation (**Fig. 6c,d, Supplementary Fig. S4a,b,c, Supplementary File S4**). Only *NveDicer1* mRNA levels displayed a mild decrease in response to the treatment (relative expression_shRNA1_ = 0.718, 95% CI, 0.470 – 1.098, p-value = 0.0966; relative expression_shRNA2_ = 0.695, 95% CI, 0.581 – 0.832, p-value = 0.0049; relative expression_shRNA3_ = 0.693, 95% CI, 0.493 – 0.972, p-value = 0.041) (**Fig. 6e, Supplementary File S4**). Altogether, our results indicate a strong link between the presence of NveRLRb and the ability to initiate downstream processes involving at least two key RNAi components i.e. NveDicer1 and NveAGO2 and a homolog of a known vertebrate IRF. Lack of effect of *NveRLRa* KD despite testing three shRNAs targeting different transcript regions together with the negative result of poly(I:C)-biotin binding assay suggest that NveRLRa might carry different functions.

**Figure 6.**
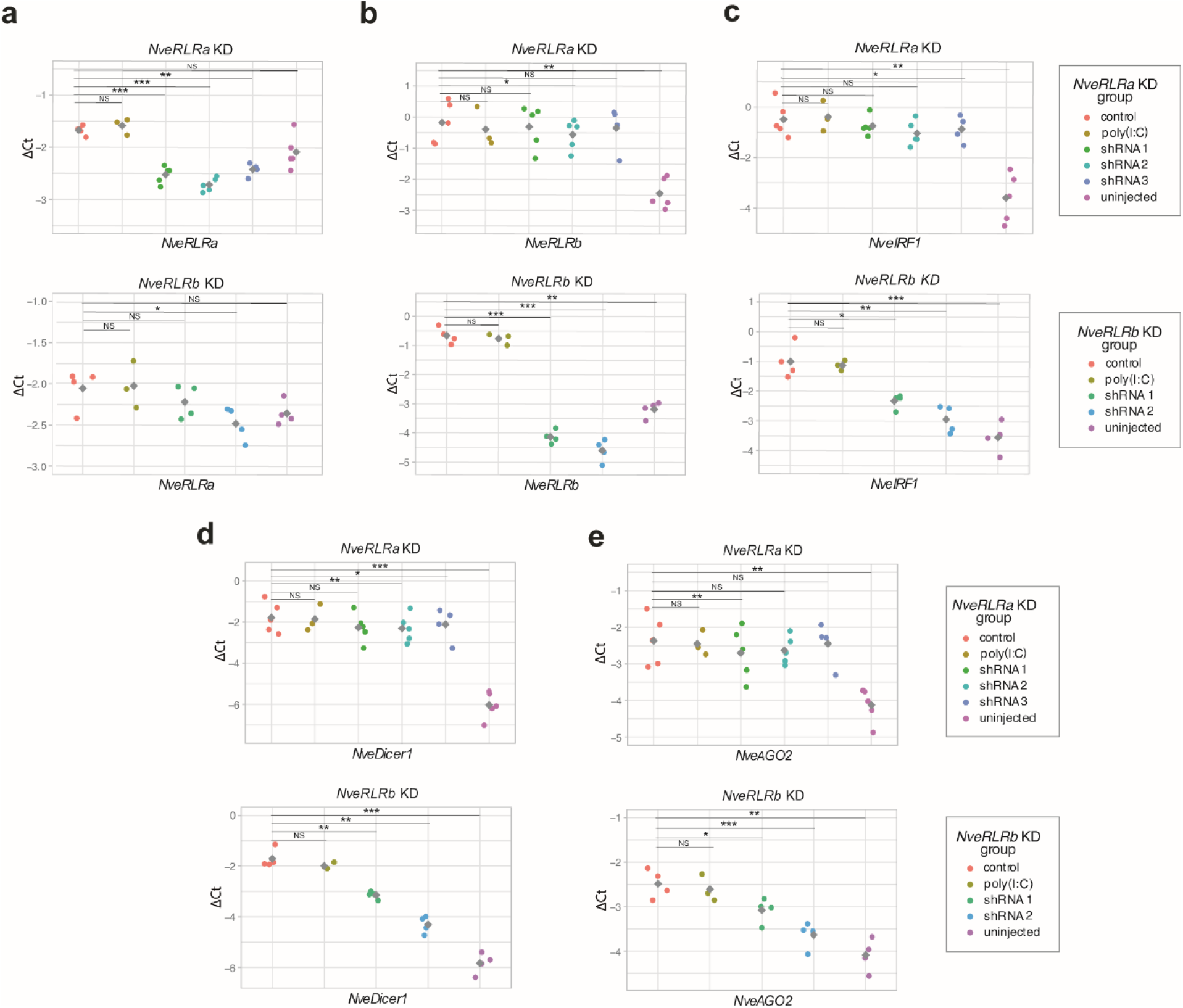
Downregulation of putative antiviral innate immunity-related genes in response to *NveRLR*s knockdown (KD) combined with poly(I:C) administration. RT-qPCR results of shRNA targeting *NveRLRa* and *NveRLRb* (upper and lower panels of each section, respectively) measuring the expression of **(a)***NveRLRa*, **(b)***NveRLRb*, **(c)***NveIRF1*, **(d)***NveDicer1*, **(e)***NveAGO2*. Grey squares represent mean values. All comparisons were done by paired two-tailed Student’s t-test against the control shRNA. Significance level: * p value < 0.05, ** p value <0.01, *** p value <0.001, NS – not significant.

## Discussion

In this study, we examined transcriptomic response to two different viral dsRNA mimics in *N. vectensis* and aimed to elucidate the role of NveRLRs in the antiviral immune pathways. We observed a lack of any signature of the antiviral immune response to the canonical RIG-I agonist which supports our hypothesis about the evolution of cnidarian RLRs from ancestral MDA5/LGP2 precursor protein **(Fig. 1a)**. In vertebrates, RIG-I binds to 5’ ends of dsRNA and recognizes the presence of di- and triphosphate on 2’-O-unmethylated nucleotide, with a strong preference to the base-paired blunt ends^9,18–22,53^. In contrast, MDA5 is known to require a stable oligomerization along the dsRNA molecule for effective downstream signaling and hence, it displays a strong affinity to long molecules with at least partial stretches of dsRNA^23–26^. Of note, poly(I:C) is known to carry 5’-diphosphate in at least a fraction of the molecules due to the synthesis process, however, uneven length of annealed strands results in single-stranded ends and long, irregular dsRNA structures^54^. In light of our results, it is likely that the activation of NveRLRs depends on the molecule length rather than the 5’ end recognition. To the best of our knowledge, the distinctive features of an effective RIG-I agonist have so far been only functionally characterized in vertebrates despite RLRs homologs being found in many invertebrate genomes. Therefore, further research on such non-vertebrate homologs can provide key insights into the evolution of dsRNA 5’ end recognition.

Transcriptomic response to poly(I:C) revealed that many canonical vertebrate antiviral factors triggered by IFN, known as interferon-stimulated genes (ISGs)^43^, are also taking part in *Nematostella* immune response. Interestingly, we observed several intriguing features of promoter region architecture such as enrichment in the TATA box sequence in poly(I:C)-upregulated genes. These elements were previously shown to display analogies in orchestrating expression of rapidly diverging and transcriptionally variable genes in phylogenetically distant groups, such as mammals^45^ and yeast^55,56^. On the other hand, response to poly(I:C) and KD experiments revealed similarities to antiviral invertebrate systems and suggested a link between NveRLRb and the RNAi pathway. Of note, a similar level of complexity and involvement of diverse antiviral mechanisms was previously suggested for the Pacific oyster *Crassostrea gigas*^57–60^, although the response to the canonical RLRs ligands presented here has not yet been comprehensively characterized in this molluscan species. The interdependence of RLRs and RNAi has been functionally demonstrated in the model nematode *Caenorhabditis elegans*, where RLRs were shown to interact with Dicer and provide crucial assistance for RNAi machinery to produce primary and secondary antiviral siRNAs^61–63^. However, unlike most other bilaterian and cnidarian RLRs, the nematode receptors lack any CARD domains (**Fig. 1b**) that typify action via oligomerization rather than association with Dicer. Importantly, there is growing evidence that virus-host interactions involve other classes of small RNAs including Dicer- and AGO-dependent microRNAs (miRNAs)^64^ and several studies in chordates suggested differential expression of host miRNAs in response to poly(I:C)^65–68^. We have recently demonstrated that the cargo of NveAGO1 is restricted to miRNAs, whereas NveAGO2 can carry both miRNAs and siRNAs^69^. This hints that poly(I:C)-upregulated NveAGO2 could function as the antiviral RNAi effector protein. Further studies will help to decipher the role of RNAi components in the antiviral immune response of *N. vectensis*.

The results of *NveRLRb* KD indicate that there are likely alternative immune cascades triggered by poly(I:C) administration which might be initiated by other dsRNA sensors. Among these, Toll-like receptors (TLRs) are obvious candidates due to their well-known role as PRRs^70^. However, the only TLR of *N. vectensis* has been recently shown to mediate immune response in NFκB-dependent way in response to *Vibrio coralliilyticus* and flagellin^71^ which indicates its involvement in recognizing bacterial rather than viral pathogens. An intriguing question for future studies is whether NveRLRa is acting as a nucleic acid sensor. On one hand the stable co-existence of two separately-clustering RLRs paralogs in sea anemones (**Fig. 1a**) and the clear increase in *NveRLRa* expression upon poly(I:C) challenge (**Fig. 4a,c**) suggest that it is likely a functional component of antiviral immune response which might display affinity to yet uncharacterized ligands. Nonetheless, the short truncation of helicase domain and aberrant KD patterns suggest an alternative but not mutually exclusive hypothesis that NveRLRa might carry some regulatory functions involved in complex feedback mechanisms.

To the best of our knowledge, our study provides the first functional insights into the role of RLRs in a non-bilaterian animal. The initial results suggest that RLRs capacity to sense 5’ end of dsRNA evolved in Bilateria, although further studies involving invertebrate RLRs will provide key answers on this matter. We show that *N. vectensis* response to a viral dsRNA mimic is characterized by high complexity and includes both vertebrate-like features, as well as invertebrate-like involvement of RNAi machinery in an RLR-dependent manner. This hints that key elements of both extant antiviral systems were already present in a cnidarian-bilaterian common ancestor. Our results lay the foundation for further functional studies on downstream effector mechanisms in *N. vectensis* which might provide key insights into the evolution of the antiviral immune response in Metazoa.

## Materials and Methods

### Sea anemone culture

*Nematostella* embryos, larvae and juveniles were grown in the dark at 22 °C in 16‰ artificial seawater, while polyps were grown at 18 °C and fed with *Artemia salina* nauplii three times a week. The induction of gamete spawning was performed as previously described^72^. The gelatinous egg sack was removed using 3% L-Cysteine (Merck Millipore, Burlington, MA, USA) and followed by microinjection of viral mimics or shRNAs.

### Injection of viral mimics

To stimulate the antiviral immune response in *Nematostella*, we used two types of synthetic dsRNA. To mimic the presence of long dsRNA, we used 3.125 ng/μl of high molecular weight (HMW) poly(I:C) in 0.9% NaCl (Invivogen, San Diego, CA, USA) with an average size of 1.5 – 8 kb, and 0.9% NaCl as a control. The second type of ligand was a synthetic dsRNA 19-mer with 5’ triphosphate (5’ppp-dsRNA) and a control dsRNA 19-mer with 5’ hydroxyl group (5’ppp-dsRNA control), both suspended in sterile RNase-free endotoxin-free water to a final concentration of 90 ng/μl (Invivogen). Each experiment was performed in triplicates and each biological replicate was composed of 100-150 injected zygotes per time point. Within each biological replicate zygotes were collected at 6, 24 and 48 hpi, flash frozen in liquid nitrogen and stored at −80 °C until processed.

### Transcriptome library preparation and sequencing

Total RNA was extracted with Tri-Reagent (Sigma-Aldrich, St. Louis, MO, USA) according to manufacturer’s protocol, treated with 2 μl of Turbo DNAse (Thermo Fisher Scientific, Waltham, MA, USA) and re-extracted with Tri-Reagent and 20 ug of RNA-grade glycogen (Thermo Fisher Scientific). The quality of total RNA was assessed on Bioanalyzer Nanochip (Agilent, Santa Clara, CA, USA) and only samples with RNA Integrity Number (RIN) > 7 were retained. Libraries were constructed from 226 ng and 300 ng of total RNA from poly(I:C) and 5’ppp-dsRNA injected samples, respectively. RNA-seq libraries were generated using SENSE Total RNA-seq Library Prep Kit v2 (Lexogen, Vienna, Austria) following the manufacturer’s protocol and sequenced on NextSeq 500 (Illumina, San Diego, CA, USA) using single-end 75 bp chemistry.

### Raw reads processing and differential gene expression analysis

Quality of raw reads was assessed and visualized with FastQC software^73^. Reads were trimmed and quality filtered by Trimmomatic with the following parameters (HEADCROP:9 LEADING:3 TRAILING:3 SLIDINGWINDOW:4:20 MINLEN:36)^74^ and the quality of the filtered reads was re-assessed in FastQC. Reads were mapped to *N. vectensis* genome (NCBI accession: GCA_000209225.1)^75^ with STAR alignment program^76^. Gene counts were obtained with RSEM^77^ (genes models, protein models and annotations are available at: https://figshare.com/articles/Nematostella_vectensis_transcriptome_and_gene_models_v2_0/807696). Differential gene expression analysis was carried out with edgeR^78^ and DESeq2^79^ implemented in the Trinity pipeline^80^. Treatment samples within each time point were compared to the corresponding control samples. Differentially expressed genes were defined by FDR < 0.05 and log2|fold change| ≥ 2. Only genes identified by both edgeR and DESeq2 methods were reported as differentially expressed. GO groups were identified by GSEA using GOseq Bioconductor package^81^ implemented in the in-built Trinity pipeline^80^. An FDR cut-off of 0.05 was considered significant for the enriched or depleted GO terms. To reduce redundancy, GO terms were group based on semantic similarity using REVIGO^82^ and visualized by CirGO v2.0^83^.

### shRNA generation and knockdown experiments

Three shRNA precursors from three different regions of each *NveRLR* gene as well as control shRNAs were designed and prepared as previously described^84^ with minor modifications. In brief, 19 bp gene targeting motif size was chosen for each shRNA (minimum GC% content > 35%). We have introduced 2-3 mismatches to the star strand, which corresponds to the coding strand, to create the bulges in shRNA precursors following the structure of native miRNA in *Nematostella*^69,85^. Reverse complement sequence of shRNA precursors was synthesized as ultramer oligo by Integrated DNA Technologies (Coralville, IA, USA), mixed with T7 promoter primer in 1:1 ratio in a final concentration of 25μM, denatured at 98 °C for 5 min and cooled to 24 °C. shRNAs were synthesized with AmpliScribe™ T7-Flash™ Transcription Kit (Epicentre, Charlotte, NC, USA) for 15 h followed by 15 min treatment with 1 μl of DNAse I. The in-vitro transcribed products were purified using the Quick-RNA Miniprep Kit (Zymo Research, Irvine, CA, USA). The quality and size of each precursor were checked on 1.5% agarose gel and its concentration was measured by spectrophotometer. The sequences of shRNAs precursors are provided in **Supplementary File S5**.

Initial screening of shRNA knockdown efficiency and toxicity revealed that microinjections of shRNAs of *NveRLRa* and *NveRLRb* proved effective and non-toxic at 48 hpi in 750-1000 ng/μl and 350-500 ng/μl concentration range, respectively. 3 shRNAs for *NveRLRa* (750 ng/μl, 750 ng/μl and 1000 ng/μl) and 2 shRNAs for *NveRLRb* (500 ng/μl each) were microinjected to *Nematostella* zygotes in a 10 μl mixture containing additionally 3.125 ng/μl of HMW poly(I:C), 1 μl of 9% NaCl and RNAse-free endotoxin-free water. Identically prepared 1000 ng/μl and 500 ng/μl of the control shRNA was included in each microinjection of *NveRLRa* and *NveRLRb* shRNAs, respectively. Moreover, in each microinjection experiment we included a subset of animals treated only with poly(I:C) 3.125 ng/μl to monitor the cytotoxic effect of shRNA control. Zygotes were collected at 48 hpi, flash frozen in liquid nitrogen and stored at −80 °C until further processed.

### Reverse-transcription quantitative PCR

To validate the results of the RNA-seq and knockdown experiments, we assayed the expression levels of several candidate immune-related genes from the mammalian RLR pathway (*NveRLRa*, *NveRLRb*, *NveOAS1*, *NveIRF1*, *NveGBP1*), RNAi pathway (*NveDicer1* and *NveAGO2*) and a representative of hexacorallian-specific gene (NVE23912) by reverse-transcription quantitative PCR (RT-qPCR) at 24 hpi (RNA-seq) or 48 hpi (knockdown experiments). 3-5 biological replicates were used to validate the results of transcriptomics and poly(I:C)-shRNAs experiments, while one biological replicate was used to assess the efficiency and background immune response to shRNAs and poly(I:C) control. RNA was extracted from injected embryos following the same protocol used for RNA-seq libraries construction. cDNA was constructed using SuperScript III (Thermo Fisher Scientific) for RNA-seq validation and iScript cDNA Synthesis Kit (Bio-Rad, Hercules, CA, USA) for knockdown experiments, according to the manufacturer’s protocol. Real-Time PCR was prepared with Fast SYBR™ Green Master Mix (Thermo Fisher Scientific) on the StepOnePlus Real-Time PCR System (ABI, Thermo Fisher Scientific). The qPCR mixture contained cDNA template (1 μl), 2× Fast SYBR™ Green Master Mix (5 μl), primers (1 μl) and nuclease-free water to make up 10 μl total volume. qPCR thermocycling conditions were 95 °C for 20 s, 40 cycles of 95 °C for 3 s, 60 °C for 30 s. Melt curve analysis was initiated with 95 °C for 15 s and performed from 60 to 95 °C in 0.5 °C increments. The expression levels of candidate genes were normalized to the NVE5273 gene (ΔCt = Ct_reference gene_ - Ct_gene of interest_) and the relative expression was calculated by the 2^ΔΔCt^ method. The significance level was calculated by applying paired two-tailed Student’s t-test to ΔCt values for each of the pairwise comparisons. Sequences of all primers are shown in **Supplementary File S5**.

### Antibody generation

For NveRLRa and NveRLRb Western blots following poly(I:C) stimulation, we used custom polyclonal antibodies raised against recombinant fragment antigens generated by rabbits’ immunization (GenScript, Piscataway Township, NJ, USA). Each recombinant fragment was injected into three rabbits. After the third round of immunization, pre-immune and post-immune sera were sent to us for screening by Western blot against *Nematostella* lysate to identify sera specifically positive for NveRLRa and NveRLRb (bands of ~111 and ~121 kDa respectively). Finally, the antigens were used by the company for affinity purification from the relevant rabbits. Amino acid sequences of NveRLRa and NveRLRb fragments used for immunization are presented in **Supplementary File S5**.

### Western blot

Equal amounts of protein were run on 4 – 15% Mini-PROTEAN® TGX™ Precast Protein Gel (Bio-Rad) followed by blotting to a Polyvinylidene fluoride (PVDF) membrane (Bio-Rad). Next, the membrane was washed with TBST buffer (20 mM Tris pH 7.6, 150 mM NaCl, 0.1% Tween 20) and blocked (5% skim milk in TBST) for 1 hour on the shaker at room temperature. Polyclonal antibody against NveRLRa or NveRLRb or monoclonal mouse anti-FLAG M2 antibody (Sigma-Aldrich) or monoclonal mouse anti-GAPDH (Abcam, Cambridge, UK) serving as loading control was diluted 1:1000 in TBST containing 5% BSA (MP Biomedicals, Irvine, CA, USA) and incubated with the membrane in a sealed sterile plastic bag at 4°C overnight. The membrane was washed three times with TBST for 10 min and incubated for 1 hour with 1:10,000 diluted peroxidase-conjugated anti-mouse or anti-rabbit antibody (Jackson ImmunoResearch, West Grove, PA, USA) in 5% skim milk in TBST. Finally, the membrane was washed three times with TBST and detection was performed with the Clarity™ Max ECL kit for pulldown experiments (Bio-Rad) and Clarity™ ECL kit for all other experiments (Bio-Rad) according to the manufacturer’s instructions and visualized with a CCD camera of the Odyssey Fc imaging system (Li-COR Biosciences, USA). Size determination was carried out by simultaneously running Precision Plus Protein™ Dual Color Protein Ladder (Bio-Rad) and scanning at 700 nm wavelength.

### Cloning and transgenesis

Synthetic genes (Gene Universal, Newark, DE, USA) including CDS of *NveRLRa* and *NveRLRb* (scaffold_15:1090025-1101489 and scaffold_40:683898-697394, respectively), self-cleaving porcine teschovirus-1 2A sequence (P2A)^86^ and mCherry sequence^87^ were amplified with Q5® Hot Start High-Fidelity DNA Polymerase (New England Biolabs, Ipswich, MA, USA), visualized on 1% agarose gel and purified by NucleoSpin Gel and PCR Clean‑up (Macherey-Nagel, Düren, Germany). Following digestion with restriction enzymes, PCR fragments were ligated to a pER242^88^ vector containing a TBP promoter previously proved to drive ubiquitous expression in *Nematostella*^89^, three N-terminal FLAG tags and SV40 polyadenylation signal. Plasmids were transformed into the *E. coli* DH5α (New England Biolabs) strain and outsourced for Sanger sequencing (HyLabs, Rehovot, Israel). Each *NveRLR* plasmid was subsequently injected into *N. vectensis* zygotes along with the yeast meganuclease I-*Sce*I (New England Biolabs) to enable genomic integration^88,90^. Transgenic animals were visualized under an SMZ18 stereomicroscope equipped with a DS-Qi2 camera (Nikon, Tokyo, Japan) and positive animals were reared to the adult stage. At approximately 4 months old F_0_ individuals were induced for gametes and crossed with wild-type animals to generate F_1_ FLAG-tagged TBP::NveRLR::mCherry heterozygotes. Positive F_1_ individuals were selected and grown to the adult stage. For the in vitro binding assay, only F_1_ females descending from a single F_0_ founder of each NveRLR line were used. Sequences of all used primers are provided in **Supplementary File S5.**

### *In vitro* binding assay

Maternal deposition of FLAG-tagged TBP::NveRLR::mCherry transgene in F_2_ animals was visualized under an SMZ18 stereomicroscope equipped with a DS-Qi2 camera (Nikon) and confirmed by Western blotting. Following fertilization with wild-type gametes, F_2_ FLAG-tagged TBP::NveRLR::mCherry and wild-type zygotes were treated with 3% L-Cysteine (Merck Millipore), washed and snap frozen in liquid nitrogen. Next, animals were mechanically homogenized in the following lysis buffer: 50 mM Tris-HCl (pH 7.4), 150 mM KCl, 0.5% NP-40, 10% glycerol, Protease inhibitor cOmplete ULTRA tablets (Roche, Basel, Switzerland) and Protease Inhibitor Cocktail Set III, EDTA-Free (Merck Millipore). Protease inhibitors were added fresh just before use. After 1 h rotation in 4°C the samples were centrifuged at 16000 × *g*, 15 min, 4 °C and supernatant was collected. Protein concentration was measured using Pierce BCA Protein Assay Kit (Thermo Fisher Scientific). Next, the lysate was pre-cleared as following: 100 μl of streptavidin magnetic beads (New England Biolabs) were washed in 1 ml of 1×PBS for 3 times and the FLAG-tagged TBP::NveRLR/wild-type lysate was added to the washed beads. Lysis buffer was added to make up 1.2 ml and samples were incubated at 4 °C rotation for 1 hour. After the incubation, the pre-cleared lysates were collected and mixed with the HMW poly(I:C) (Invivogen) or HMW poly(I:C)-biotin (Invivogen) in the final concentration of 30 ng/ml and ATP (New England Biolabs) in the final concentration of 0.5 mM. Samples were incubated for 1 h in rotation at room temperature. Simultaneously, 100 μl of fresh streptavidin magnetic beads were blocked with wild-type lysates alike in the pre-clearing step. poly(I:C) samples were added to the blocked beads and incubated for 2 h in rotation at 4 °C for poly(I:C)-biotin pulldown. After the incubation, the lysates were discarded and the beads were washed 3 times with 500 μl of the following wash buffer: 50 mM Tris-HCl (pH 7.4), Protease inhibitor cOmplete ULTRA tablets (Roche) and Protease Inhibitor Cocktail Set III, EDTA-Free (Merck Millipore). Subsequently, 40 μl of filtered double-distilled water and 20 μl of Blue Protein Loading Dye (New England Biolabs) were added to the beads. The samples were heated at 100 °C for 8 min and placed on ice for 1 min, then centrifuged 1 min at 21,000 × *g* at 4 °C, and the supernatant was collected for Western blot.

### Phylogenetic analysis

To construct an informative phylogenetic tree we selected representatives of major groups carrying RLRs: vertebrates (a fish, an amphibian, and a mammal), two non-vertebrate chordates (a lancelet and a lamprey), nematodes (*C. elegans* and *A. suum*), two lophotrochozoans (an oyster and an annelid) and hexacorallians (three sea anemones, each representing a different major sea anemone clade and two-reef building corals). Sponges RLRs sequences were chosen as an outgroup. The RLRs amino acid sequences were aligned using MUSCLE^91^ and low certainty alignment regions were removed by TrimAl^92^ using the –automatic1 for heuristic model selection. The maximum-likelihood phylogenetic trees were constructed using IQ-Tree^93^ with the LG+F+R5 model which was the best fitting model both according to the Bayesian information criterion (BIC) and corrected Akaike information criterion. Support values of the ML tree were calculated using 1,000 ultrafast bootstrap replicates^94^. A Bayesian tree was constructed using MrBayes^95^ with the WAG +I +G model and the run lasted 5,000,000 generations with every 100th generation being sampled. The Bayesian analysis was estimated to reach convergence when the potential scale reduction factor (PSRF) reached 1.0. Consensus domain composition was predicted by simultaneous search in Pfam^96^ and NCBI Conserved Domains^97^ databases run with default parameters.

Homologs of NVE23912 sequences were identified through a search in TSA and NCBI nr databases and *Nematostella* gene models. Amino acid sequences were aligned using MUSCLE^91^ and visualized by CLC Genomics Workbench. Details of RLRs and NVE23912 homolog sequences used in the analysis are available in **Supplementary File S6**.

### Promoter sequence analysis of DEG

Analysis of promoter sequences was performed as previously described^45^ with minor modifications. In brief, coordinates of the TSS were retrieved from nveGenes.vienna130208.nemVec1.bed file. We subset the upregulated DEG identified by poly(I:C) microinjection (n=1379) and the fraction of top 10% genes (n=138) and top 20% genes (n=276), setting the whole transcriptome as the background (n=18831). TATA box-containing genes were identified using FIMO^98^ by having at least one statistically significant match (p-value cut-off of <0.05) to the TATA box consensus motif (MA0108.1) retrieved from JASPAR server^99^. Due to uncertainty in TSS calling, we have scanned the coding strand in two ways: a) narrow search included 38 bp upstream of TSS; b) wide search spanned both 100 bp upstream and 100 bp downstream of putative TSS whenever fitted in the scaffold boundaries. To estimate motifs enrichment in the same groups, we used the non-redundant JASPAR core motif matrix (pfm_vertebrates.txt) and run AME^100^ in one-tailed Fisher’s exact test mode. The searching region included 500 bp upstream of the putative TSS, the first exon and the first intron of the gene. For motif identification, the cut-off of adjusted by Bonferroni correction p-value < 0.05 was considered significant statistically significant. The presence of the signal peptide in each protein sequence was predicted by SignalP 4.1 Server with default settings^101^.

### Data availability

All sequencing data that support the findings of this study have been deposited in the National Center for Biotechnology Information Sequence Read Archive (SRA) and are accessible through the BioProject accession number PRJNA673983. Source data for Fig. 2,3c and Supplementary Fig. 1a,b,c,d,e have been provided in Supplementary File S1. Source data for Fig. 4a,b, 6 and Supplementary Fig. 6 have been provided in Supplementary File S4. All other relevant data are available from the corresponding authors on request.

