## Supplementary Figures for "Functional characterization of the cnidarian antiviral immune response reveals ancestral complexity"

**Content:**

**Supplementary Figure. S1:** Differential gene expression among control and viral mimics.

**Supplementary Figure S2:** Alignment of NVE23912-like homologs identified in Hexacorallia species.

**Supplementary Figure S3:** Frequency of genes with predicted TATA box elements and signal peptide within poly(I:C)-induced genes.

**Supplementary Figure S4:** Response of putative antiviral innate immunity-related genes to poly(I:C) and *NveRLR*s knockdown (KD) combined with poly(I:C).

**Supplementary Figure S5:** Background immune response to shRNA and controls for viral mimics.

**Supplementary Figure S6:** NveRLRs protein level in various developmental stages.

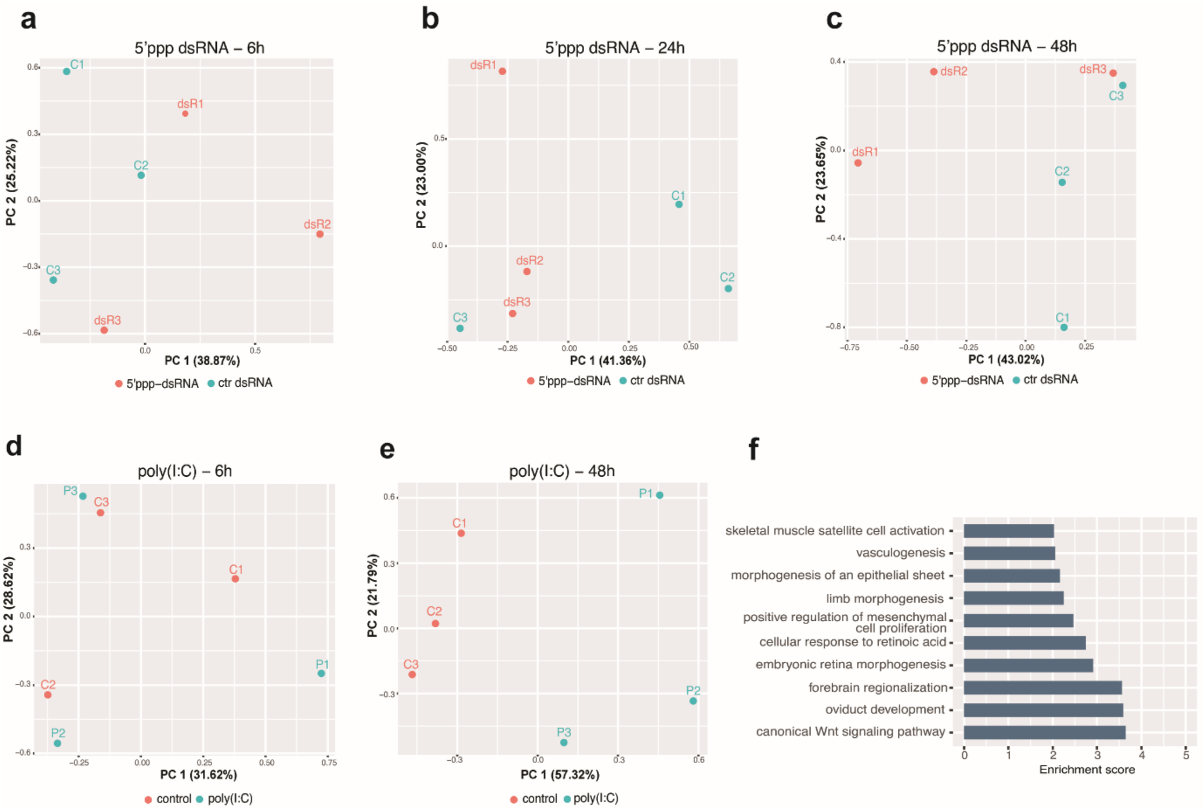


**Figure S1. Differential gene expression among control and viral mimics.** PCA plots representing whole transcriptome of **(a-c)** short 5'ppp dsRNA-injected animals and **(d-e)** poly(I:C)-injected animals assayed at different time points. **(f)** GO terms enrichment after REVIGO-based semantic similarity filtering of downregulated DEG at 6 hpi after short 5'ppp dsRNA injection. Enrichment score was defined as -($log_{10}$ p value).

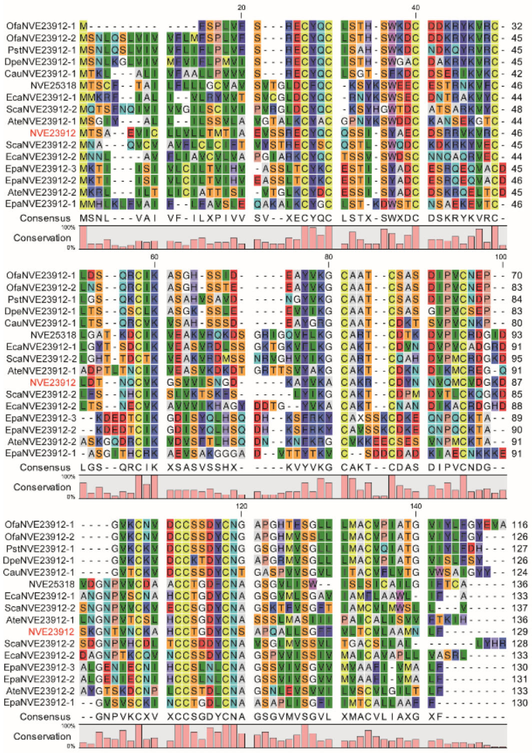

**Figure S2. Alignment of NVE23912-like homologs identified in Hexacorallia species.** Ate, *Actinia tenebrosa*, Cau, *Corynactis australis*, Dpe, *Desmophyllum pertusum* (previously *Lophelia pertusa*), Eca, *Edwardsiella carnea*, Epa, *Exaiptasia pallida*, Nve, *Nematostella vectensis*, Ofa, *Orbicella faveolate*, Pst, *Pseudodiploria strigosa*, Sca, *Scolanthus callimorphus*.

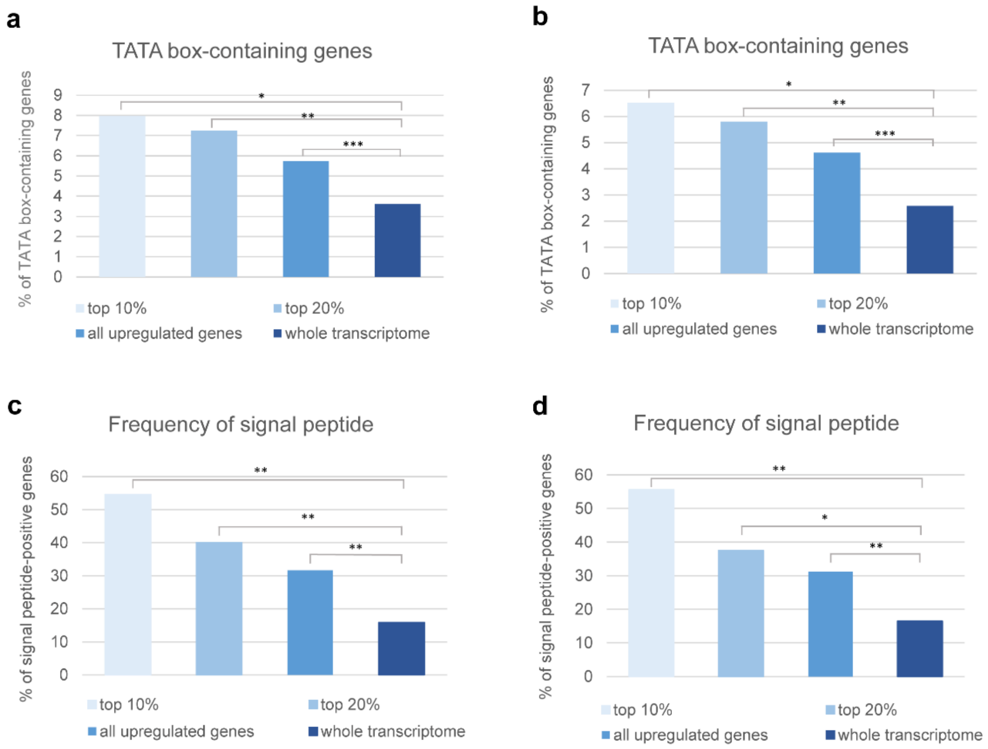


**Figure S3. Frequency of genes with predicted TATA box elements and signal peptide within poly(I:C)-induced genes.** Frequency of TATA-box positive genes were identified by search within **(a)** 100 bp upstream and 100 bp downstream of TSS, and **(b)** 38 bp upstream of TSS. Signal peptide was identified in the same **(c)** wide and **(d)** narrow search windows. Significance level was assessed by two-tailed Fisher's exact test; * p value < 0.05, ** p value <0.01, *** p value <0.001.

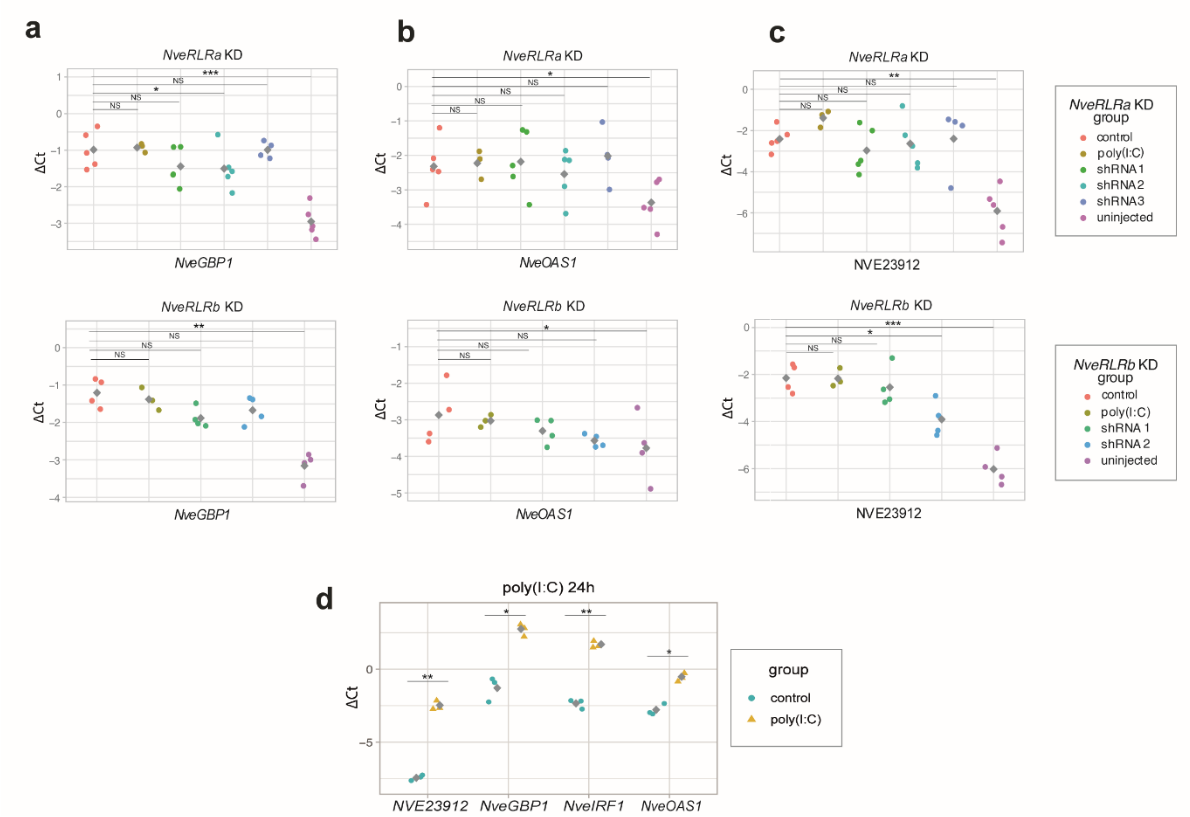


**Figure S4. Response of putative antiviral innate immunity-related genes to poly(I:C) and *NveRLRs* knockdown (KD) combined with poly(I:C).** RT-qPCR results of multiple shRNA targeting *NveRLRa* and *NveRLRb* (upper and lower panels of each section, respectively) assaying expression of **(a)** *NveGBP1*, **(b)** *NveOAS1*, **(c)** NVE23912. **(d)** Upregulation of putative antiviral innate immunity-related genes in response to poly(I:C) injection after 24 h. Grey squares represent mean values. All comparisons were done by paired two-tailed Student's t-test against the control shRNA or 0.9% NaCl. Significance level: * p value < 0.05, ** p value <0.01, *** p value <0.001.

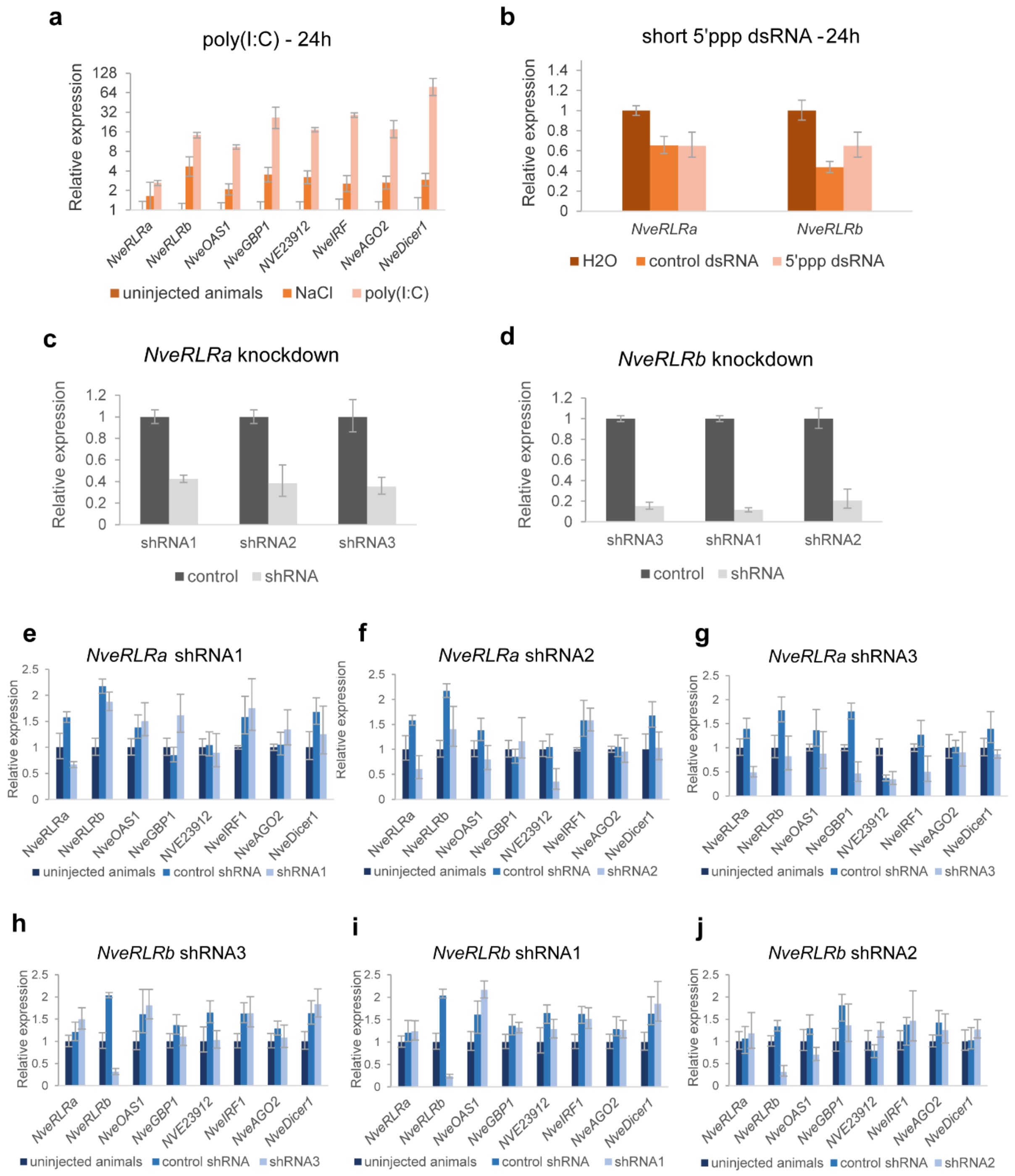


**Figure S5. Background immune response to shRNA and controls for viral mimics.** Relative gene expression level of selected putative immune-related genes in response to **(a)** poly(I:C) and 0.9% NaCl, **(b)** short dsRNA with and without 5' triphosphate. Knockdown efficiency of **(c)** NveRLRa shRNAs and **(d)** NveRLRb shRNAs. Expression of putative immune-related genes in response to **(e-g)** *NveRLRa* shRNAs, **(h-j)** *NveRLRb* shRNAs. Error bars represent standard deviation of technical replicates.

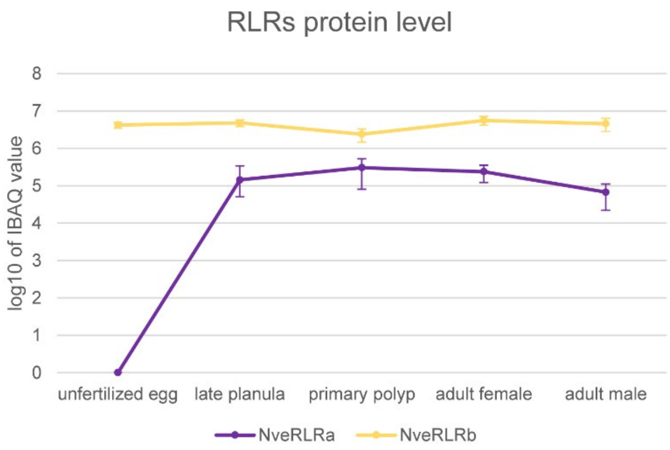


**Figure S6. NveRLRs protein level in various developmental stages.** A graph presents at a logarithmic scale the mass spectrometry-measured iBAQ (Intensity Based Absolute Quantification) values. Error bars represent standard deviation. Data acquired from Columbus-Shenkar *et al.*[1]
